# Genomic insights into ancestry and infectious disease in 17th-century colonial Brazil

**DOI:** 10.64898/2025.12.10.693243

**Authors:** Ksenia Macias Calix, Caroline Borges, Ana Lucia do Nascimento Oliveira, Suely Cristina Albuquerque de Luna, Claudia Minervina Souza Cunha, Alexsandra Maria de Siqueira, Andre Luiz Campelo dos Santos, Michael DeGiorgio, John Lindo, Raquel Assis

## Abstract

During urban redevelopment in the historic district of a Brazilian metropolis, archaeologists uncovered a previously undocumented 17th-century cemetery, containing the closely spaced remains of over two dozen young adult males of military age. Historical records suggest the site once housed a makeshift hospital, raising questions about the origins and causes of death of those interred, particularly given the absence of skeletal evidence for violent or fatal trauma. The current study integrates bioarchaeological, historical, and genomic data to investigate the ancestry and putative disease-related mortality of nine individuals whose remains were recovered and subsequently sequenced. Population-genetic analyses revealed strong affinities with Northern European populations, particularly from Norway, Iceland, Estonia, and Czechoslovia, consistent with their likely roles as soldiers or laborers employed by the Dutch West India Company. To explore potential causes of death, we conducted a metagenomic screening with a novel pipeline optimized for degraded DNA, which revealed widespread presence of *Klebsiella pneumoniae* and *Mycobacterium tuberculosis* pathogens across all samples. Authenticity was confirmed through post-mortem damage patterns characteristic of historical samples. These findings, together with the absence of combat trauma and the collective burial context at the site, support the hypothesis of an epidemic-related mortality event. This study contributes to the growing field of historical pathogen genomics and offers a rare genomic perspective on life, mobility, and health during a period of colonial upheaval in South America.

## Introduction

Between 1630 and 1654, a large portion of colonial Brazil came under Dutch control during a period known as Dutch Brazil or New Holland [Boxer, 1957]. Governed by the Dutch West India Company, the occupation was centered in the territory of the current state of Pernambuco and encompassed the major cities of Recife and Olinda, the current and former state capitals. This region held immense strategic and economic value due to its sugarcane plantations and access to Atlantic trade networks. Under the leadership of Johan Maurits of Nassau (1637–1644), Dutch Brazil became a hub of not only military and commercial activity but also scientific exploration and cultural exchange. Artists, naturalists, and cartographers documented the landscape and peoples of the colony, providing one of the earliest European records of Brazil’s social and ecological landscapes [Boxer, 1957, Miranda, 2020]. The period was marked by intense interactions among Indigenous communities, enslaved Africans, and European colonists, leaving lasting imprints on the region’s demographic and epidemiological history [Meuwese, 2012]. This colonial backdrop is essential for interpreting archaeological findings from 17th-century Brazil, particularly those involving burial contexts linked to sites of military or medical activity, where multiple cultural and biological influences may converge.

An intriguing discovery in 2014 brought new attention to the colonial past of Recife, when construction workers in the historic district uncovered a previously unknown cemetery beneath the foundations of centuries-old buildings. This unexpected finding occurred during urban redevelopment in the community of Pilar, and hereafter the archaeological site will be referred to as the Pilar cemetery. Excavations in the area revealed dozens of human burials with no prior record of a cemetery at that location. The location of the Pilar cemetery within the State of Pernambuco, Brazil, is shown in Figure 1A. Systematic excavations later revealed that the burial area extended across several adjacent city blocks and contained more than 130 individuals [Silva, 2015, da Silva Oliveira, 2024]. Stratigraphic evidence indicated that the deepest burials corresponded to the earliest phase of use, dated to the late sixteenth and early seventeenth centuries [Silva et al., 2019]. Spatial patterns varied across excavation sectors: some blocks, such as Quadra 55, displayed orderly interments oriented toward the coastline, whereas others contained shallow, irregular, or commingled burials, consistent with rapid deposition during periods of crisis. These archaeological observations help contextualize the demographic uniformity and mortuary characteristics of the individuals analyzed in this study and frame the historical circumstances under which the cemetery operated. Historical maps, however, indicated the area once housed the Sao Jorge Fortress, which may have functioned as a field hospital during Dutch occupation [Silva, 2015]. The combination of military infrastructure and public health concerns during the Dutch occupation makes this a historically plausible context for the formation of a burial ground, particularly one that may have operated during times of conflict or epidemic.

**Figure 1:**
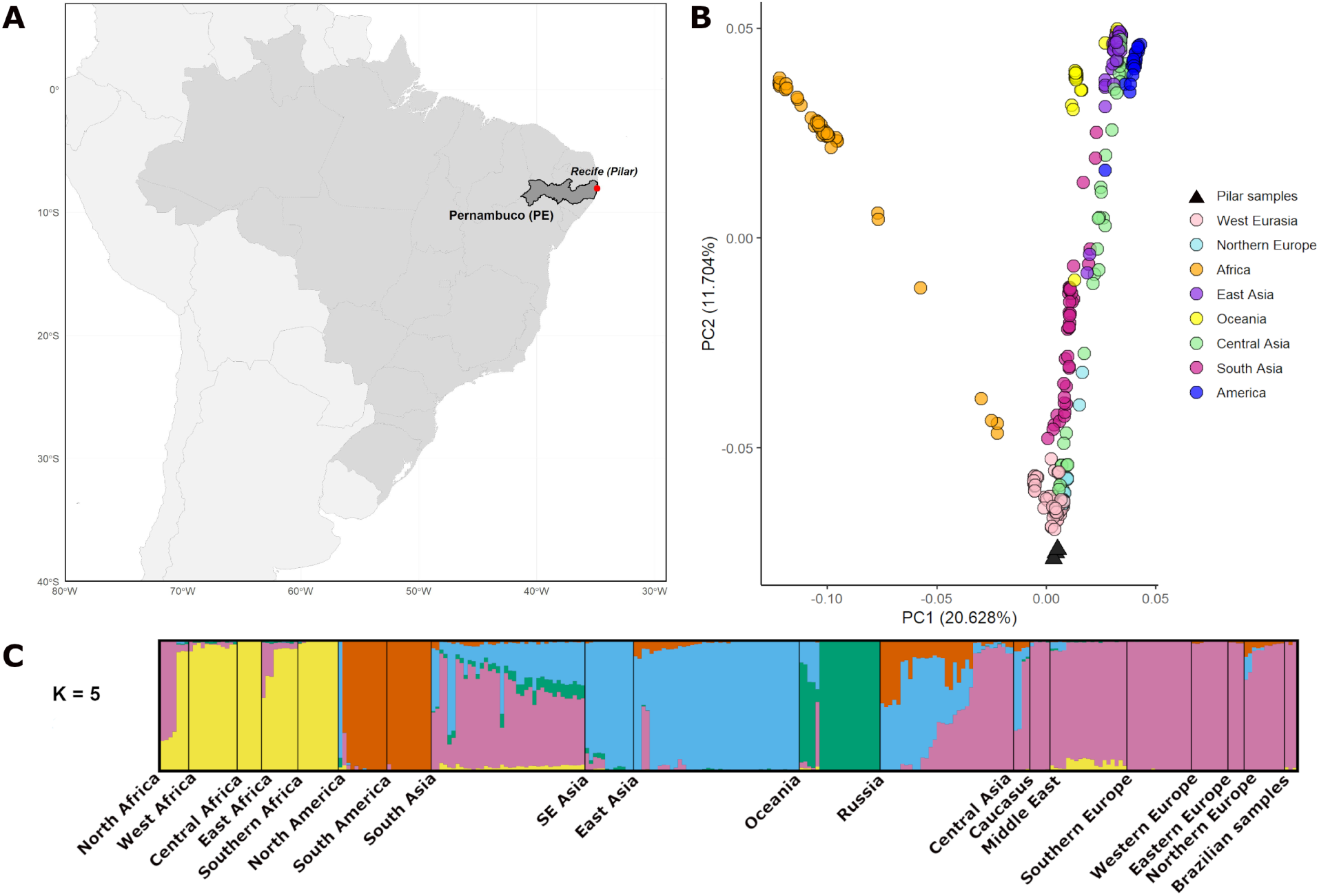
Overview of site location and ancestry analyses for the Pilar individuals. (A) Geographic map showing the location of the Pilar archaeological site within Recife, Brazil. (B) PCA of the three highest-coverage Pilar individuals (black triangles), projected onto present-day reference populations from the SGDP dataset. Reference population colors correspond to major geographic regions. (C) ADMIXTURE analysis results. Each vertical bar corresponds to one individual and is subdivided into up to five colored ancestry components, with segment height indicating the proportion of each component. The three highest-coverage individuals are shown in the final segment of the plot. Expanded results for all nine Pilar individuals are provided in supplementary Figures S2 (PCA) and S3 (ADMIXTURE).

Bioarchaeological analyses of the remains provided initial insights into the demographic profiles of the buried individuals. All skeletons recovered from this sector of the site belonged to young adult males, generally aged between 15 and 21 at time of death. Based on non-genetic assessments, Silva [2015] proposed that these individuals were likely of European origin; however, such evaluations based solely on osteological observations, should be regarded as provisional, particularly in colonial contexts characterized by migration and admixture. We therefore treated these interpretations as hypotheses to be evaluated using genomic data. The uniformity in sex and age, together with the cemetery’s location and its apparently limited period of use, suggests that the burial ground may have served a specific social group, possibly military recruits or laborers associated with the Dutch West India Company. Genetic analyses thus play a central role in clarifying the ancestry of these individuals and refining our understanding of their place within the broader demographic landscape of Dutch Brazil.

Beyond origins, the cause of death for this homogeneous group remains a pressing question. Further analysis of burial practices at the site strengthened the hypothesis of a rapid, large-scale mortality event. At least six graves contained multiple individuals, often buried without apparent pattern or grave goods, and in positions suggesting hasty interment [Silva, 2015]. No weapons, military gear, or personal items were recovered, and none of the skeletons showed evidence of perimortem trauma or battle wounds, making it unlikely these individuals perished in combat. Instead, the absence of funerary organization, coupled with the consistent age and sex profile of the individuals, points toward a sudden health crisis, such as an epidemic. Archaeological indicators of physiological stress support this interpretation. Several individuals displayed skeletal signs of scurvy, including alveolar bone resorption, as well as periosteal reactions and healed trauma indicative of chronic malnutrition and systemic stress [Silva, 2015]. These conditions would have compromised immune function and increased vulnerability to infectious diseases. Historical records from Dutch Recife describe recurring outbreaks of illness during the occupation, exacerbated by poor water quality, overcrowding, and irregular food supplies [Melo, 2002, Miranda, 2014, Ujvari, 2003]. Epidemic episodes—including dysentery, scurvy, verminoses, and yellow fever—were reportedly common, with one outbreak in 1646 said to have overwhelmed the Sao Jorge hospital and caused the deaths of hundreds of colonists [Silva, 2011].

In this context, the convergence of bioarchaeological and historical evidence supports the hypothesis that these individuals died during a localized disease outbreak. However, historical documentation provides limited medical detail, and osteological signs alone are rarely pathogen-specific. To address these limitations, this study combines population-genetic and metagenomic approaches with archaeological and historical context to investigate the ancestry and causes of death of nine individuals recovered from the Pilar cemetery. We extracted DNA from skeletal remains and applied population-genetic methods to assess whether these individuals shared ancestry with European groups, as previously hypothesized based on preliminary morphological assessment. In parallel, we conducted metagenomic screening using a novel pipeline to identify microbial taxa and evaluate the presence of pathogenic infections. Together, these methods allowed us to reconstruct both the ancestral origins and the potential biological contributors to the sudden deaths of individuals recovered from the Pilar cemetery. This integrated design illustrates the value of genomic analyses guided by historical and archaeological frameworks for illuminating identity and health in early colonial contexts.

## Results

The first question we sought to address was the ancestry of the individuals in the Pilar cemetery. As an initial step, we assessed maternal and paternal lineages using mitochondrial DNA haplogroup assignment and Y-chromosome haplogroup inference (Table S1; see *Methods*). All nine individuals were assigned to West Eurasian mitochondrial clades commonly observed in present-day European populations. Y-chromosome haplogroups were successfully inferred for all individuals, most of whom carried paternal lineages within R1a or R1b, which are both widely distributed across Europe. One individual (IND23A) carried a haplogroup primarily associated with East African populations, and another (IND12) could only be assigned to an upstream clade that did not allow clear regional attribution. Taken together, these uniparental lineages are broadly consistent with predominantly European ancestry among the Pilar individuals and provide contextual support for subsequent autosomal analyses.

As is typical for ancient DNA datasets, genome-wide coverage varied markedly across individuals (Figure S1, Table S1). Therefore, while we performed all downstream analyses on the nine individuals, we focus our interpretations on the three highest-coverage genomes (IND21, IND36B, and IND51A), which retained sufficient SNP density for robust inference. Results for the remaining individuals are included in Figures S2–S12, and are interpreted with appropriate caution given their lower coverage.

To investigate genome-wide ancestry patterns, we performed principal component analysis (PCA) [Patterson et al., 2006], using a heterogenous set of present-day reference populations from the Simons Genome Diversity Project (SGDP) [Mallick et al., 2016] spanning multiple global regions (see *Methods*). The PCA results show that our samples project most closely to populations from West Eurasia and Northern Europe (Figure 1B), providing strong support for their proposed European origin. PCA results for all nine individuals are shown in Figure S2. As expected, lower-coverage genomes display more variable projections, with one individual appearing shifted toward the area of principal-component space associated with African reference groups. We interpret these shifts as reflecting projection instability due to missing data rather than genuine affinity differences. Such behavior is well documented in PCA projections of low-coverage ancient genomes [Zabel et al., 2025, François and Jay, 2020] and does not indicate African ancestry. Importantly, the overall pattern remains consistent, with the majority of samples falling within the West Eurasian range. Although preliminary, these findings align with the archaeological conclusions, offering initial evidence of a genetic affinity between Pilar individuals and populations from Europe.

To further characterize ancestry and refine the resolution of these findings, we conducted an ADMIXTURE analysis [Alexander et al., 2009]. The optimal number of ancestral components that best explained the data was determined to be *K* = 5 through cross-validation (see *Methods*). Visualization of these results reinforced our PCA findings, indicating that the primary ancestral component of the three best-preserved Pilar individuals is overwhelmingly shared with present-day populations from Europe, encompassing regions of Western, Northern, Southern, and Eastern Europe (Figure 1C). This same component also shows notable representation in the Caucasus and, to a lesser extent, in parts of South Asia. ADMIXTURE results across values from *K* = 2 to 10 for all nine individuals are shown in Figure S3, and display the same overall pattern, with the strongest shared ancestry traced to European populations. Overall, the ADMIXTURE results provide additional support for the strong European affinities of the Pilar individuals and align with the broader historical context of Dutch colonial settlement in Brazil.

To build on our ancestry findings, we computed outgroup *f*_3_ statistics [Patterson et al., 2012], which estimate genetic affinities by measuring shared genetic drift between two populations relative to an outgroup population. For each calculation, we used one of our highest-quality samples (IND21, IND36B, or IND51A) as a reference and computed outgroup *f*_3_ statistics against each of 65 present-day comparison populations from the SGDP panel, using the Yoruba population as the outgroup (see *Methods*). The outgroup *f*_3_ results supported our previous findings, revealing strongest genetic affinities between the Pilar individuals and populations from Northern Europe and West Eurasia (Figure 2). IND21 showed highest affinities with Iceland, Norway, Estonia, Czechoslovakia, and Poland, while IND36B displayed a nearly identical pattern, with top affinities to Estonia, Czechoslovakia, Iceland, Orkney Islands, and Poland. IND51A likewise clustered most closely with Norway, Iceland, Czechoslovakia, France, and the Orkney Islands. These patterns were further supported by *D*-statistic tests, which showed the strongest allele sharing between the Pilar individuals and populations from Northern and Western Europe. The results of outgroup *f*_3_ and *D*-statistic tests for all nine individuals are presented in Figures S4-S12 and show a broadly similar affinity profile. Together, these results reinforce our earlier analyses (Figure 1), highlighting consistent genomic associations between our samples and populations from Northern and Western Europe, while providing a finer resolution of their ancestral structure.

**Figure 2:**
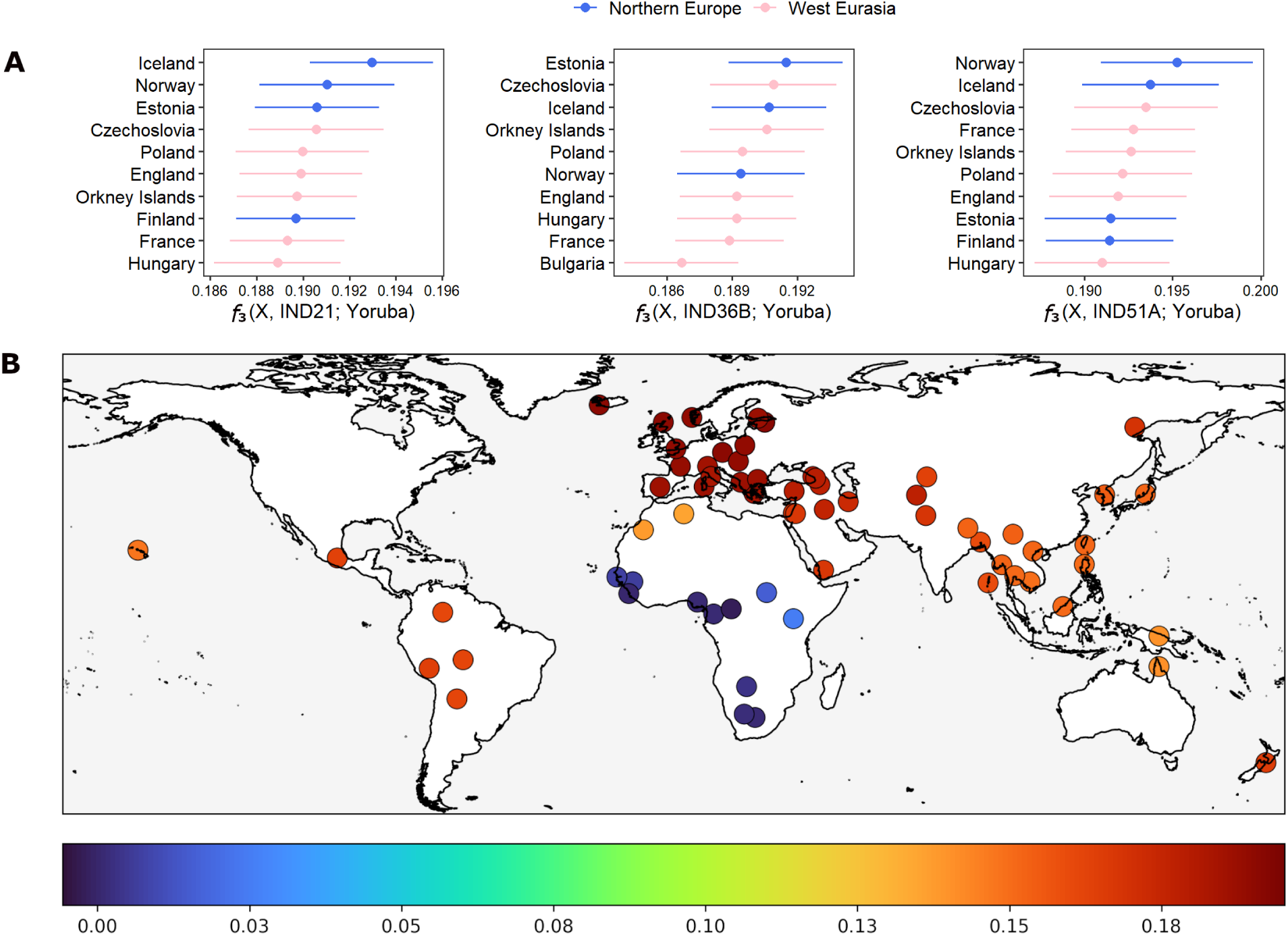
Outgroup *f*_3_ analyses summarizing genetic affinities between the Pilar individuals and present-day reference populations. (A) Outgroup *f*_3_ statistics ranked by decreasing value, depicting genetic affinity between the three best-preserved Pilar individuals IND21, IND36B, and IND51A, shown from left to right. The *x*-axis represents *f*_3_ scores, where higher values indicate greater genetic affinity or shared ancestry, and error bars denote the standard errors of the *f*_3_ estimates. (B) Geographical visualization of outgroup *f*_3_ affinities, where affinity strength is represented by a color gradient ranging from blue (lowest *f*_3_ values) to red (highest *f*_3_ values). Because the affinity patterns of all three individuals were nearly identical in geographic distribution and magnitude, a single representative map based on individual with highest coverage (IND36B) is shown.

Next, we investigated the potential role of an epidemic event as a cause of death of the individuals in the Pilar cemetery. To address this question, we developed a targeted metagenomic screening pipeline with safeguards tailored to ancient DNA (Figure 3A). We first aligned all sequencing reads to the human reference genome and extracted the unmapped reads to enrich for non-human sequences and maximize efficiency in downstream microbial detection. These reads were analyzed with Kaiju [Menzel et al., 2016], a protein-level classifier that offers enhanced sensitivity for detecting the degraded and divergent sequences typical of ancient samples. For this study, we applied Kaiju with its most comprehensive pathogenic reference database, enabling broad taxonomic identification and enhancing our ability to detect rare or low-abundance pathogens. To further refine results, we filtered Kaiju assignments through the NMDC GCPathogen database [Guo et al., 2024], retaining only human-specific pathogens and ranking taxa by abundance in the best-preserved samples.

**Figure 3:**
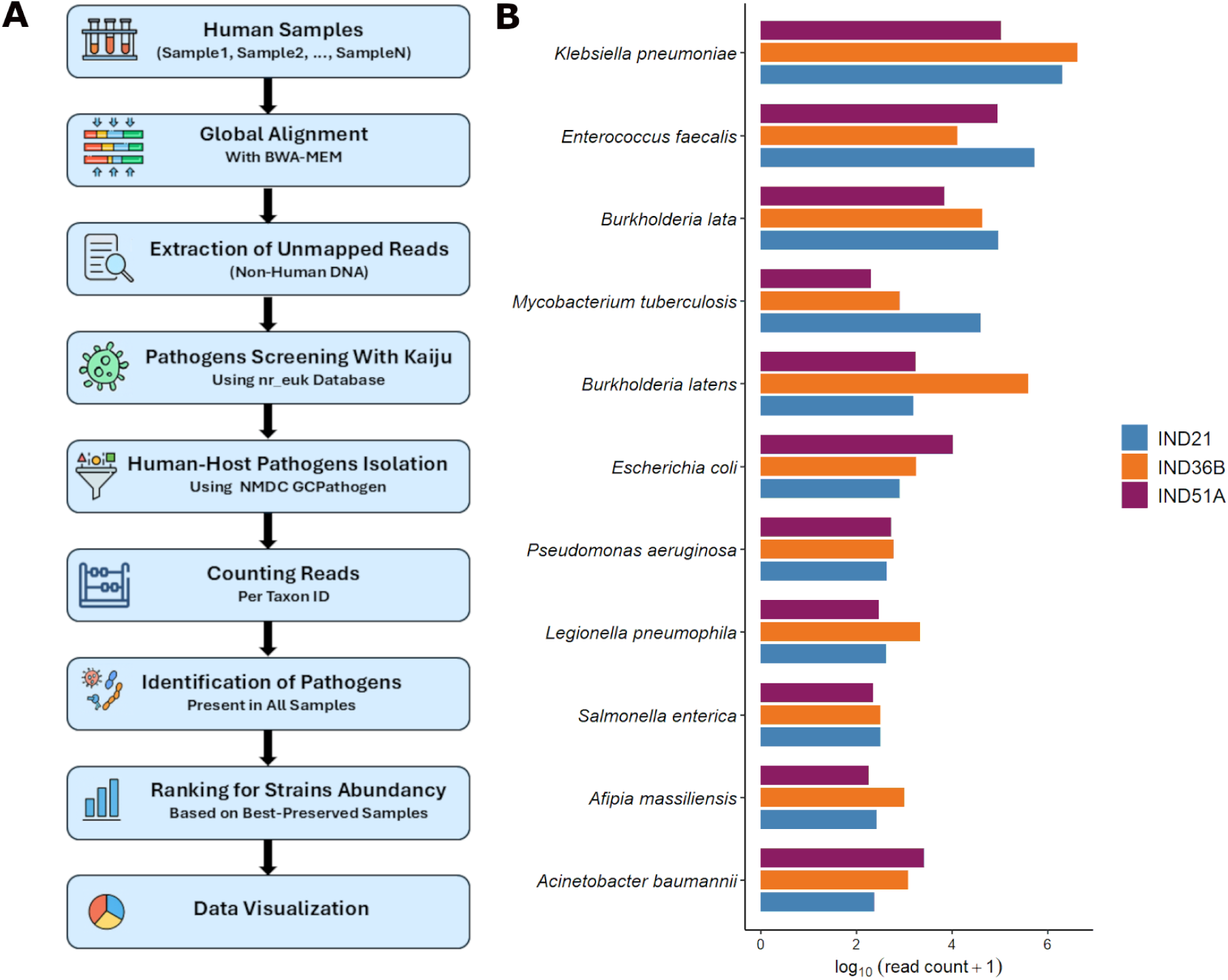
Overview of the metagenomic screening workflow and pathogen detection results. (A) Pipeline developed for pathogen detection, including reads alignment, filtering, taxonomic classification, and authentication steps (see *Methods*). (B) Barplots showing the log-transformed read counts of the top pathogenic taxa identified in the best-presernved samples IND21, IND36B, and IND51A.

This pipeline identified a range of microbial taxa across the dataset (Figure 3B). Some species, such as *Burkholderia lata*, *Burkholderia latens*, and *Legionella pneumophila*, are commonly associated with environmental sources like soil and water [Hall et al., 2015, Heijnsbergen et al., 2014], while others, including *Escherichia coli* and *Salmonella enterica*, are part of the normal gut flora [Marian et al., 2024, Martinson and Walk, 2020]. These taxa likely represent environmental background organisms or postmortem microbial colonizers, which commonly appear in ancient remains [Hofreiter et al., 2001, Warinner et al., 2017]. In contrast, several clinically relevant taxa were consistently detected across all nine individuals. Most notably, *Klebsiella pneumoniae* and *Mycobacterium tuberculosis* emerged as dominant candidates. In our three best-preserved samples (IND21, IND36B and IND51A), we identified 2,019,174, 4,157,465, and 107,047 reads classified as *K. pneumoniae*, and 40,307, 801, and 203 reads classified as *M. tuberculosis*, respectively. Opportunistic pathogens, such as *Enterococcus faecalis* and *Acinetobacter baumannii* [Freitas et al., 2016, Howard et al., 2012], were also present, consistent with immune suppression and possible nosocomial transmission [Nithichanon et al., 2022, Peleg et al., 2008]. To assess authenticity, we evaluated post-mortem DNA damage [Skoglund et al., 2014], which confirmed characteristically elevated cytosine deamination patterns in both human and microbial sequences, particularly for *M. tuberculosis* and *K. pneumoniae* (Figure S13). Together, these findings support the hypothesis that these two pathogens contributed to the deaths of the individuals recovered from the Pilar cemetery.

## Discussion

The genetic findings and archaeological context presented in this study provide compelling evidence that the individuals buried at the previously undocumented Pilar cemetery were of Northern European origin. Specifically, our PCA, ADMIXTURE analysis, outgroup *f*_3_ statistics, and *D*-statistics consistently revealed genomic affinity between the Pilar samples and European populations. Notably, the three highest-quality ancient genomes showed particularly strong genetic affinities with Northern European populations, especially those from Iceland, Norway, and Estonia. These results support earlier non-genetic hypotheses of European ancestry of the Pilar individuals, specifically estimating Northern Europe as their most likely region of origin.

Our metagenomic screening, performed using a pipeline tailored for detecting microbial DNA in highly degraded and low-coverage samples, revealed substantial traces of *K. pneumoniae* and *M. tuberculosis* across all individuals, with the highest read counts in the three best-preserved samples. Characteristic ancient DNA damage patterns confirmed the authenticity of both human and microbial sequences, supporting their origin from degraded historical material [Skoglund et al., 2014]. The co-occurrence of these two pathogens, both known to cause pulmonary and systemic infections, and to spread rapidly in overcrowded, malnourished populations [Cioboata et al., 2025, Ockenga et al., 2023, Sahu et al., 2024, Sinha et al., 2019], offers a compelling explanation for the abrupt, large-scale mortality event suggested by the archaeological context. Taken together with the demographic profile of the deceased, the disorderly nature of their interment, and spatial characteristics of the burial site [Silva, 2015], our findings support the interpretation that this cemetery functioned as an emergency burial ground during an epidemic that struck a cohort of young European men stationed far from home.

Despite the significance of these findings, several limitations should be acknowledged. While DNA was successfully extracted from nine individuals and all exhibited consistent patterns in both ancestry and pathogen presence, only three samples yielded sufficient endogenous content to allow for detailed and statistically robust analyses. Although the depth of interpretation is necessarily greater for these three higher-quality genomes, the broader consistency observed across the entire sample set reinforces the reliability of our conclusions and strengthens the overall narrative derived from this cohort. Second, the sequencing protocols were optimized for human DNA, as the original study design focused on questions of ancestry. Consequently, while we detected reliable signals from *M. tuberculosis* and *K. pneumoniae*, the relatively low microbial coverage precluded genome assembly, strain typing, or phylogenetic reconstruction of the pathogens. Nevertheless, the ability to detect ancient pathogenic DNA from human-focused sequencing libraries demonstrates the resilience of paleogenomic data and underscores the potential for future pathogen discovery even from sub-optimally targeted datasets [Spyrou et al., 2019].

This study offers an important contribution to the growing field of historical pathogen genomics, particularly within the understudied context of colonial South America. By combining genetic data with archaeological and historical evidence, it provides new insights into the identities, health conditions, and causes of death among a previously undocumented population, while also highlighting the potential of interdisciplinary approaches to shed light on past epidemic events. The implications of this work extend well beyond the historical period it investigates. By applying paleogenomic tools to a little-studied colonial context in South America, this study highlights the potential for reconstructing pathogen presence and spread in early modern populations. It contributes to ongoing work aimed at integrating pathogen detection into ancient genomic research [Andrades Valtueña et al., 2017] and offers a valuable reference point for future comparative studies of immune response and disease susceptibility across historical and modern genomes.

Building on these findings, future research could explore how genetic variation in immune-related genes has influenced human susceptibility to respiratory pathogens over time. Comparative studies between historical populations and modern genomic datasets offer a promising avenue to investigate long-term patterns of host-pathogen co-evolution, particularly in genes involved in innate and adaptive immunity, such as *TLR2*, *NOD2*, *IL6*, and *IFNGR1*, which have been previously implicated in differential responses to mycobacterial and opportunistic infections [Hu and Spaink, 2022, Joseph and Lindo, 2022, Landes et al., 2015]. Such research could help clarify whether past exposure to pathogens like *K. pneumoniae* and *M. tuberculosis* has influenced the evolution of human immune genes in ways that are still observable in the genomes of people living today. Integrating historical genomic data with contemporary epidemiological patterns may also shed light on enduring vulnerabilities or resistances in human populations, offering a deeper evolutionary perspective on infectious disease dynamics [Barreiro and Quintana-Murci, 2010, Bos et al., 2011].

In conclusion, this study opens a rare window into the demographic and epidemiological landscape of 17th-century Brazil, shedding light on a little-known population affected by disease during a period of colonial upheaval. By combining genomic analysis with archaeological and historical context, we were able to reconstruct aspects of origin, health, and mortality in individuals who left few traces in written records. Beyond its immediate findings, this work demonstrates the broader value of applying molecular tools to historical human remains, particularly in regions and time periods that remain underexplored in genomic research. As genomic methods continue to advance, such interdisciplinary efforts will be essential for tracing how migration, environment, and infectious disease have jointly shaped human biological diversity. Ultimately, this study underscores how genetic traces preserved in historical remains can help us better understand the forces that continue to shape human identity and health today.

## Methods

### Archaeological context and sample recovery

The skeletal remains analyzed in this study were recovered during salvage excavations carried out in 2014 in the historic district of Recife, Pernambuco, Brazil. These excavations revealed a previously undocumented 17th-century burial area, now referred to as the Pilar cemetery [Silva, 2015, da Silva Oliveira, 2024]. Systematic stratigraphic work across several adjacent city blocks identified more than 130 individuals interred in contexts characteristic of rapid, minimally furnished colonial-period burials.

All human remains were documented using standard archaeological procedures, including stratigraphic recording, spatial mapping, and controlled sediment removal [Balme and Paterson, 2014, Buikstra and Ubelaker, 1994]. The nine individuals included in this study were sampled based on the availability of preserved teeth suitable for ancient DNA analysis, using established guidelines for the selection and handling of skeletal material in bioarchaeological and aDNA research [Buikstra and Ubelaker, 1994, Hofreiter et al., 2001]. After archaeological recovery, remains were handled under contamination-minimizing conditions and transported using conservation and chain-of-custody practices appropriate for fragile human material [Sease, 1994].

### DNA extraction and sequencing

DNA was extracted from isolated teeth belonging to nine individuals. The nine teeth were selected based on a macroscopic/visual analysis attesting their external integrity and, therefore, high potential for endogenous DNA preservation. The extraction of ancient DNA and preparation of sequencing libraries were conducted at the Lindo Ancient DNA Laboratory at Emory University. DNA was extracted from approximately 50–100 mg of tooth powder using a silica-based method optimized for highly degraded samples, following the protocol described by Dabney et al. [2013]. The protocol includes a decalcification step with EDTA and proteinase K digestion, followed by silica column binding in a guanidine-based buffer, multiple wash steps, and final elution in TE buffer.

Single-stranded DNA libraries were constructed using the SRSLY PicoPlus Uracil+ Library Preparation Kit (ClaretBio), which is specifically designed to retain uracil residues characteristic of ancient DNA damage patterns (Figure S13). The protocol includes denaturation, 3’ and 5’ adapter ligation, and a uracil-tolerant polymerase-based amplification step. Dual indexed libraries were pooled and sequenced on two lanes of the Illumina NovaSeq X platform at Psomagen (Rockville, MD), using 100 bp paired-end chemistry.

### Processing of genetic data

Following sequencing, raw reads were processed to remove Illumina adapters using AdapterRemoval2 [Schubert et al., 2016], ensuring high-quality read trimming. The trimmed sequences were subsequently aligned to the human reference genome (hg19) using the BWA *mem* algorithm [Li and Durbin, 2009], which provides high mapping accuracy for ancient DNA fragments [Xu et al., 2020]. Quality control metrics were assessed with MultiQC [Ewels et al., 2016], and low-quality or duplicate reads were removed prior to downstream analyses. Mean genomic depths across the nine individuals were uniformly low, with all samples falling below 2.1*×* coverage (Table S1). Such values are typical for degraded historical material and require analytical strategies designed for sparse data. Given these limitations, we used ANGSD [Korneliussen et al., 2014] to derive pseudo-haploid genotypes from BAM files by leveraging genotype likelihoods, a strategy well suited for low-coverage DNA [Gennaro et al., 2025, Bergström et al., 2022].

To establish a robust framework for population genetic analysis, we used the SGDP reference dataset [Mallick et al., 2016], which includes 278 present-day individuals representing a varied array of populations from Africa, Europe, Asia, Oceania, and the Americas. This dataset provides one of the most comprehensive resources for capturing worldwide human genomic diversity, allowing reliable comparison of ancient and modern population structure. The raw dataset contained 49,791,567 SNPs and was filtered to retain only biallelic sites with no missing data, resulting in a high-quality dataset of 28,592,222 SNPs. To minimize linkage disequilibrium and retain independent informative loci, we performed pruning across 200-kilobase windows with an *r*^2^ [Hill et al., 1966] threshold of 0.2, yielding 9,489,542 SNPs. We then applied a minor allele frequency filter of 0.05 to exclude rare variants that may reflect sequencing noise or population-specific drift, producing a final, balanced reference panel of 405,558 SNPs suitable for downstream analyses. These filtered positions were then used to guide pseudo-haploid genotype extraction from the ancient BAM files. As part of the pseudo-haploid sampling strategy, ANGSD first estimates genotype likelihoods from BAM files using the SAMtools model (-GL 1) and then uses these likelihoods to generate pseudo-haploid calls via the -doHaploCall module [Korneliussen et al., 2014]. We restricted our analysis to the positions retained in the filtered SGDP reference panel and applied stringent quality filters, including minimum mapping quality of 30 and base quality of 30, to minimize sequencing errors. Pseudo-haploid genotypes were generated by random sampling of a single high-quality base per site (-doHaploCall 1), while specifying the major and minor alleles from the reference panel (-doMajorMinor 3). The final datasets contained 190,700 SNPs for IND21, 235,132 SNPs for IND36B, and 59,049 SNPs for IND51A. For completeness, SNP counts for the remaining six individuals were as follows: 6,311 (IND12), 6,564 (IND20), 18,178 (IND23A), 8,243 (IND31B), 24,242 (IND73C), and 9,396 (IND78). These data were converted to EIGENSTRAT format using the EIGENSOFT v7.2.1 suite [Patterson et al., 2006], producing SNP, IND, and GENO files for downstream analyses.

Pseudo-haploid datasets were generated for all nine Pilar individuals and merged with the filtered SGDP reference panel to create unified datasets for downstream analyses. Using ANGSD’s likelihood-informed pseudo-haploid calling reduces potential biases introduced during variant calling and yields more consistent genotype representations across individuals and analyses, enabling a clearer assessment of population affinities despite the low coverage of the ancient genomes.

### Uniparental haplogroup inference

Mitochondrial DNA haplogroups were assigned using HaploGrep 3 (v3.2.1) [Schönherr et al., 2023] based on mtDNA reads extracted from the mapped BAM files. Y-chromosome haplogroups were inferred using Yleaf [Ralf et al., 2018], which assigns paternal lineages directly from read-level data. These analyses were used to characterize maternal and paternal ancestry components prior to genome-wide population genetic analyses.

### Principal component analysis

To assess broad genomic affinities between our samples and populations from reference datasets, and to identify potential outliers in the reference panel, we performed PCA using the *smartpca* program from the EIGENSOFT v7.2.1 package [Patterson et al., 2006]. Principal components (PCs) were computed using present-day populations specified via the *poplistname* option. Pilar samples, characterized by a high proportion of missing data, were subsequently projected onto these precomputed PCs using the *lsqproject: YES* option. Automatic shrinkage correction for eigenvalues was enabled with the *autoshrink: YES* option. We visualized the PCA results using the ggplot2 package [Wickham, 2016] in R [R Core Team, 2023], focusing on the first two principal components.

### Admixture analysis

We used ADMIXTURE v1.3.0 [Alexander et al., 2009] to investigate population structure in our dataset. This software models each individual as having ancestry from a user-defined number of clusters *K*, estimating the proportion of their genome derived from each cluster. To determine the optimal number of clusters, we performed ADMIXTURE runs with *K* ranging from 2 to 10. The optimal value for *K* was identified based on the smallest 10-fold cross-validation error after 100 iterations, using the -cv=10 and -C 100 settings. The ADMIXTURE results for all tested values of *K* were visualized using pong v1.5 [Behr et al., 2016].

### Outgroup *f*_3_ analysis

To assess affinities between the Pilar individuals and present-day populations, we performed outgroup *f*_3_ analyses [Patterson et al., 2012] using the merged dataset of the ancient samples and the SGDP reference panel [Mallick et al., 2016] (see *Processing of genetic data*). This analysis estimates shared genetic drift between two populations relative to an outgroup, providing a measure of their genetic affinity. Yoruba was chosen as the outgroup population, enabling estimation of shared genetic drift between each ancient genome and present-day reference populations. The *f*_3_ statistics were computed in the form *f*_3_(Reference1, Reference2; Yoruba) using the qp3Pop program from ADMIXTOOLS [Patterson et al., 2012]. The number of SNPs included in the calculations varied among population trios, depending on coverage and overlap with the reference panel. Across all tests, the minimum SNP count was 43, 902 for the trio (IND51A, Nigeria; Yoruba), while the maximum was 232, 972 for (IND36B, Russia; Yoruba). For the remaining six lower-coverage individuals, SNP counts ranged from 4,689 for (IND12, Sierra Leone; Yoruba) to 24,011 for (IND73C, Russia; Yoruba).

### *D*-statistics

*D*-statistic tests were conducted using the qpDstat module from ADMIXTOOLS [Patterson et al., 2012] to quantify allele-sharing asymmetries. Statistics were computed in the form *D*(*X,* Karitiana; Pilar individual, Yoruba), where *X* denotes each of the 52 reference populations tested for every individual. Yoruba was used as the outgroup, and Karitiana was included as one of the two comparison populations. This configuration allowed us to evaluate whether each Pilar individual was equidistant to population *X* and Karitiana, as expected under a tree-like relationship. To match the conditions of the outgroup *f*_3_ analysis, we used the same SGDP reference panel [Mallick et al., 2016], excluding African populations from *X* to avoid violating assumed tree relationships. In total, 468 population quartets were analyzed across the nine Pilar individuals, corresponding to tests of all 52 reference populations paired with Karitiana and Yoruba. The number of SNPs contributing to each test varied depending on overlap among the four populations. Among the three-best-preserved individuals, SNP counts ranged from 59,048 for IND51A to 235,131 for IND36B, whereas for the remaining six individuals values ranged from 6,311 for IND12 to 24,242 for IND73C. The resulting *D*-statistics were visualized using the ggplot2 R package [Wickham, 2016, R Core Team, 2023].

### Pathogen detection analysis

To determine whether a past epidemic event contributed to the deaths of the Pilar individuals, we first filtered out human-derived reads by mapping the raw sequencing data to the human reference genome using BWA *mem* [Li and Durbin, 2009] and retaining only unmapped reads with SAMtools [Li et al., 2009] *−f* 4 setting. These non-human reads were then analyzed with Kaiju (version 1.10.1) [Menzel et al., 2016], a taxonomic classification tool designed for high-throughput sequencing data from whole-genome metagenomic DNA. Kaiju performs taxonomic assignments by directly comparing sequencing reads against a reference database of microbial and viral protein sequences, utilizing the NCBI taxonomy. To maximize taxonomic coverage and sensitivity, we used the nr euk database [Menzel et al., 2016], the most comprehensive reference available. This database includes protein sequences from archaea, bacteria, viruses, fungi, and microbial eukaryotes, allowing for thorough screening of potential pathogens. To further refine our results, we leveraged the gcPathogen resource [Guo et al., 2024] to obtain a curated list of taxonomic IDs for host-specific human pathogens. This list was subsequently used to filter our Kaiju output, ensuring that only pathogens capable of infecting humans were retained for further analyses.

### Post-mortem damage analysis

To evaluate the authenticity of the ancient DNA and confirm the presence of post-mortem damage patterns characteristic of degraded genetic material, we used PMDtools v0.60 [Skoglund et al., 2014], which is designed to quantify and extract reads exhibiting deamination-induced misincorporations, primarily cytosine to thymine (C*→*T) and guanine to adenine (G*→*A) substitutions at read termini. Trimmed reads were used for all post-mortem damage analyses. For human DNA, trimmed reads were aligned to the human reference genome using BWA *mem* [Li and Durbin, 2009] under default parameters. For microbial taxa, we first extracted the read identifiers corresponding to *M. tuberculosis* and *K. pneumoniae* from the Kaiju classification output (see *Pathogen detection analysis*). These identifiers were then used to retrieve the corresponding reads from the paired-end fastq files using seqtk v1.3 [Heng Li, 2013]. The extracted reads were aligned using BWA *mem* to the *M. tuberculosis* NCBI RefSeq assembly GCF 000195955.2 and *K. pneumoniae* NCBI RefSeq assembly GCF 000240185.1. The resulting BAM files for both host and microbial alignments were input to PMDtools in deamination detection mode to produce tabular output describing terminal substitution frequencies. Visualization of post-mortem damage patterns was performed in R [R Core Team, 2024] using ggplot2 [Wickham, 2016], enabling inspection of damage profiles for each taxon. These results served to validate the ancient origin of both human and microbial DNA in our dataset.

## Data availability

Raw sequencing reads (FASTQ format) and associated metadata for the nine Pilar individuals have been deposited in the NCBI Sequence Read Archive (SRA) under BioProject accession PRJNA1374090 and BioSample accessions SAMN53670052–SAMN53670060. Data are under embargo and will be made publicly available upon publication of this article.

## Acknowledgments

This work was supported by National Institutes of Health grants R35GM142438 and R35GM128590, and by National Science Foundation grants DEB-2302258 and DBI-2130666.

## Notes

### Competing Interest Statement

The authors have declared no competing interest.

