## Supplementary material for "Genomic insights into ancestry and infectious disease in 17th-century colonial Brazil": Figures S1-S13, Table S1

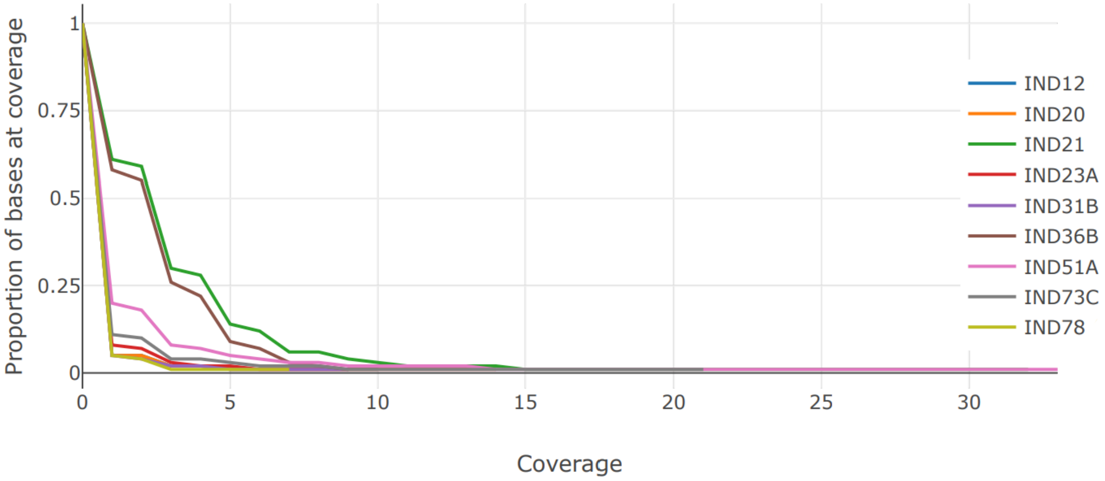


Figure S1: Coverage distribution across the nine Pilar individuals. The plot shows the proportion of genomic bases observed at each coverage level, providing an overview of sequencing depth and data completeness for all samples. Coverage values are based on genome-wide estimates.

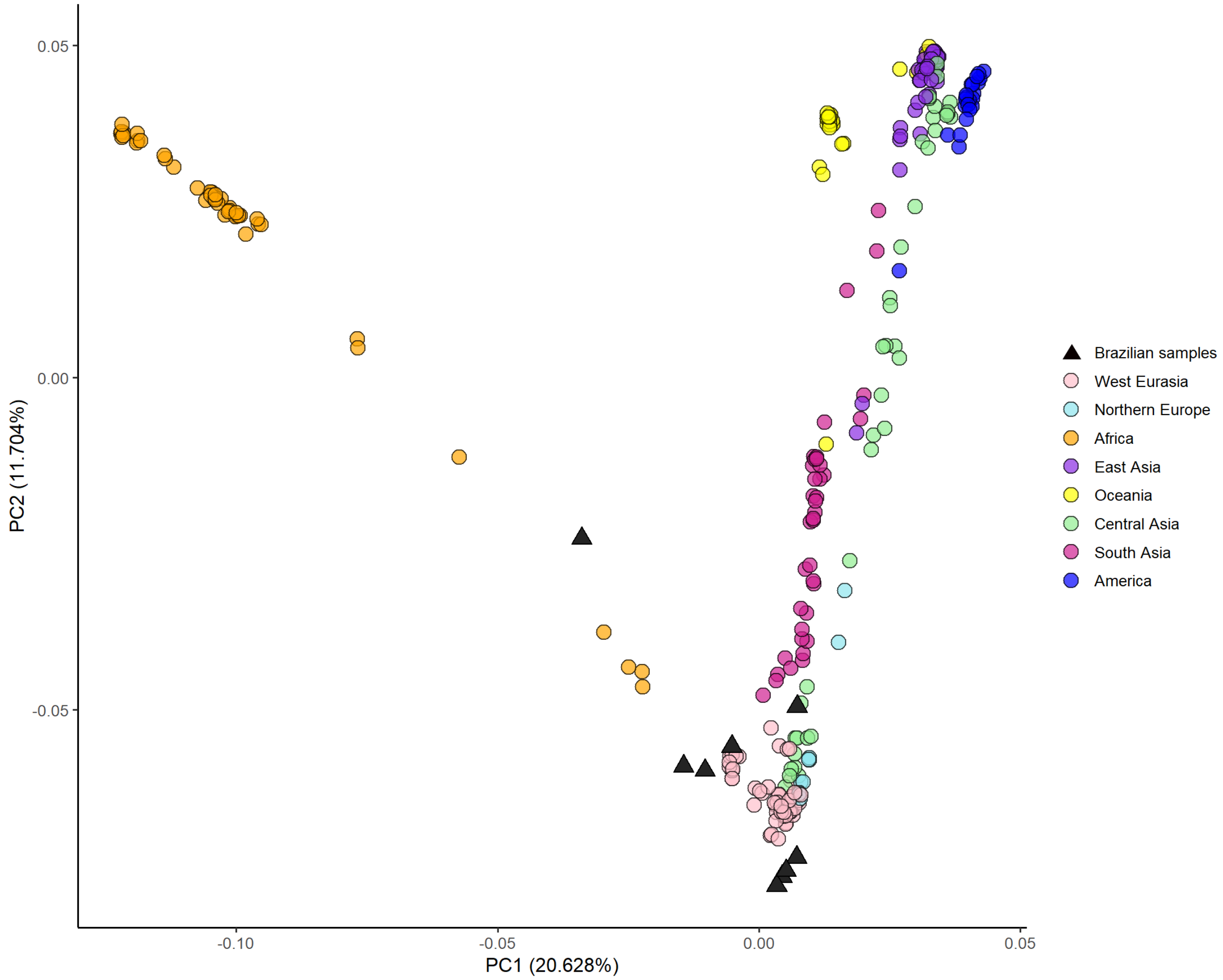


Figure S2: PCA for all nine Pilar individuals (black triangles), projected onto present-day reference populations from the SGDP dataset. Lower-coverage genomes display greater projection variability. Reference population colors correspond to major geographic regions.

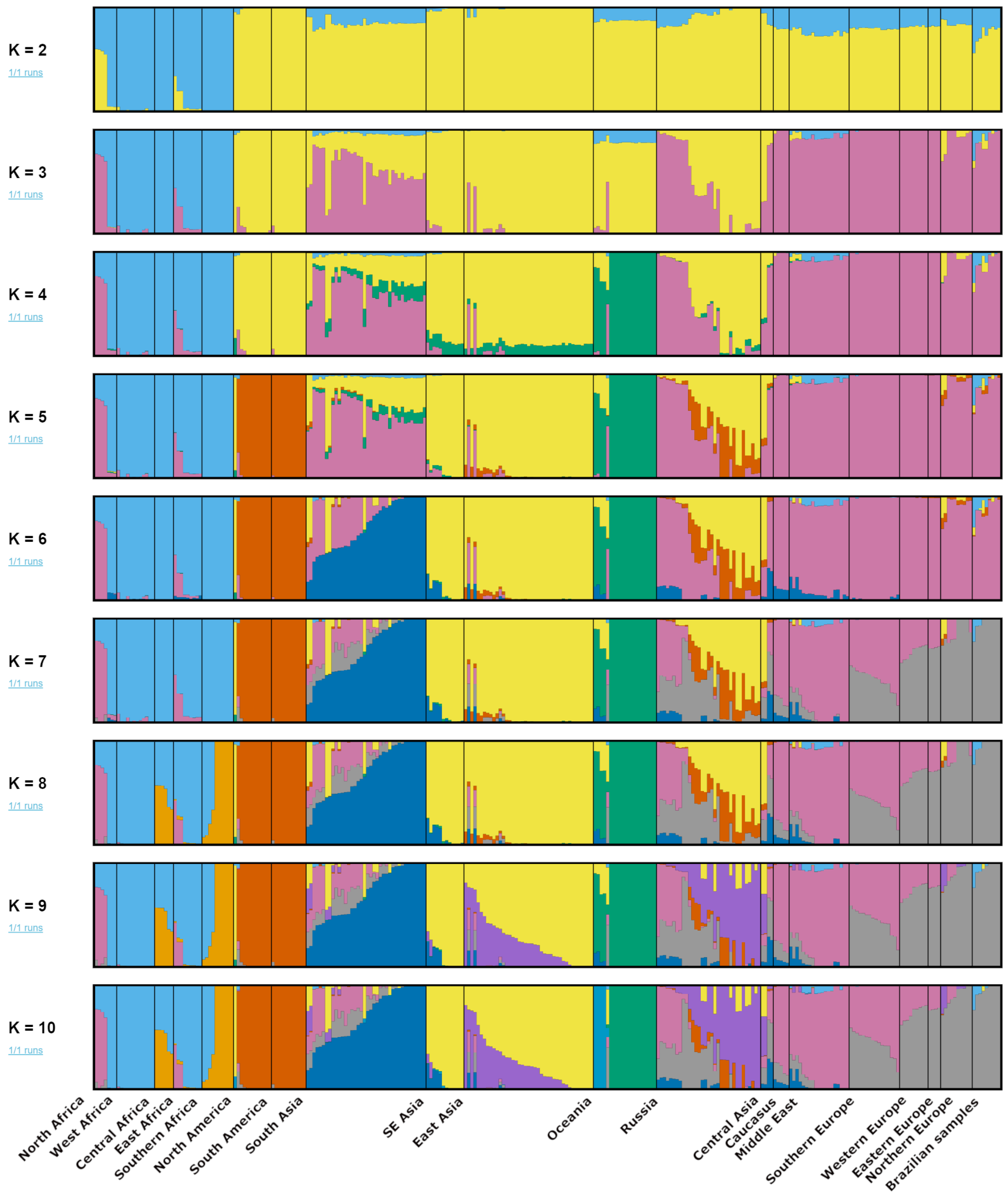


Figure S3: ADMIXTURE analysis results for $K = 2$ through $K = 10$, including all nine samples. Each vertical bar corresponds to one individual and is subdivided into ancestry components, with color segment height indicating the proportion of each component. The Pilar individuals are shown in the final segment of the plot.

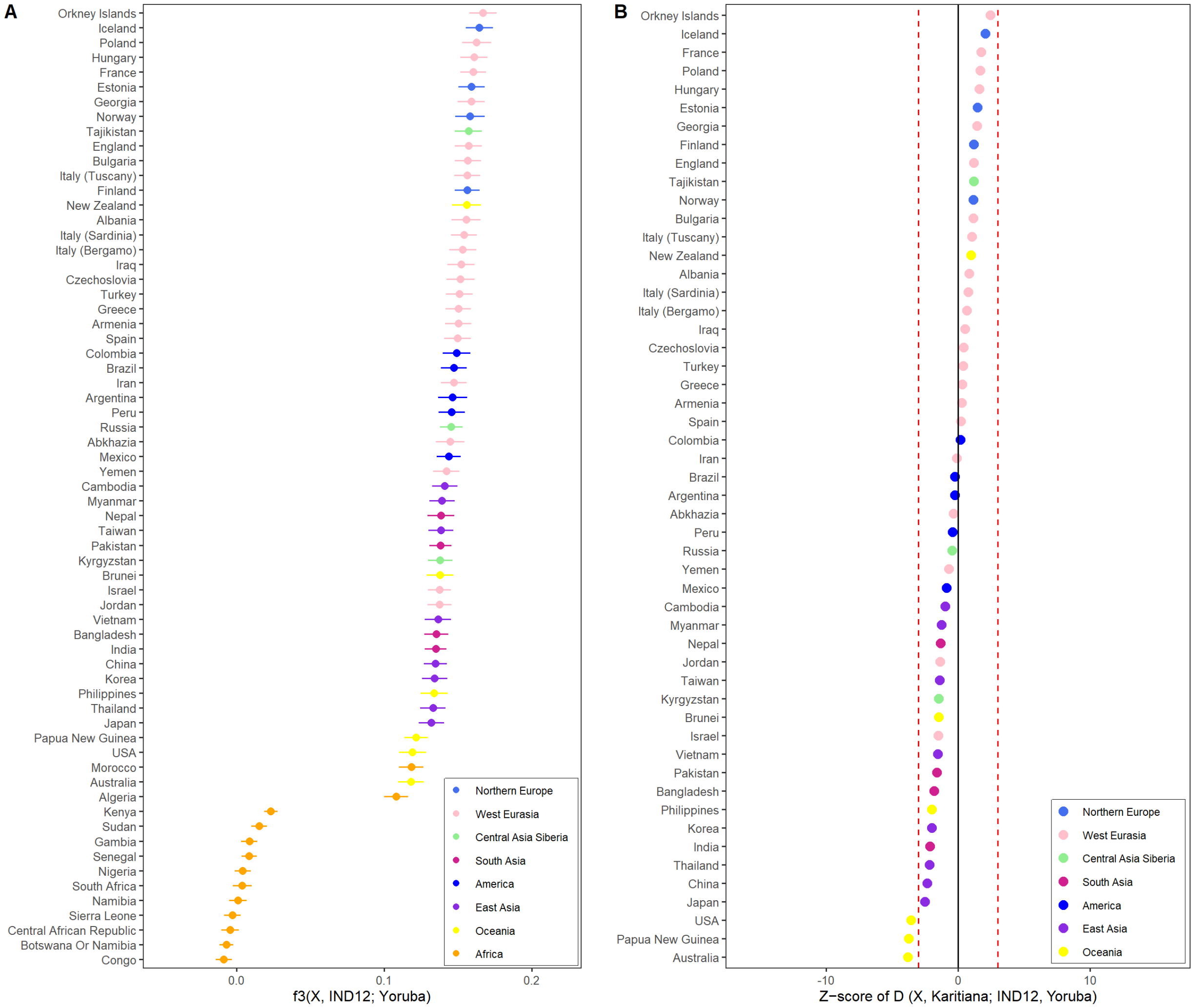


Figure S4: Outgroup $f_3$ and $D$-statistic results for IND12. (A) Outgroup $f_3$ statistics for IND12, computed in the form $f_3(X, \text{IND12}; \text{Yoruba})$ and ranked by decreasing affinity across comparison populations. (B) $D$-statistics, computed as $D(X, \text{Karitiana}; \text{IND12}, \text{Yoruba})$ and ranked by decreasing $Z$-score. Positive values indicate excess allele sharing between IND12 and population $X$ relative to Karitiana, while negative values indicate the reverse. Vertical dashed lines at $Z = \pm 3$ mark the conventional thresholds used to assess significance.

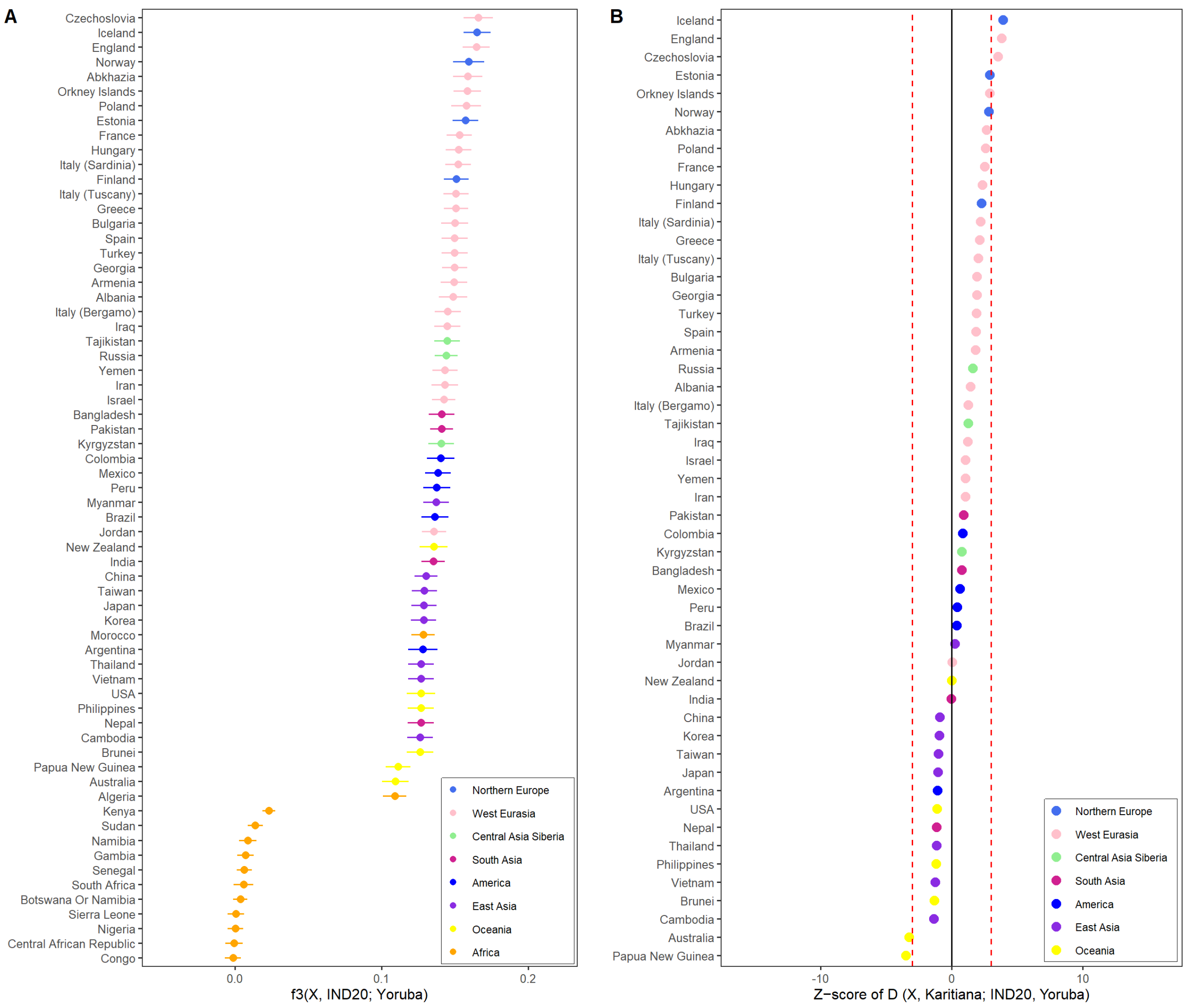


Figure S5: Outgroup $f_3$ and $D$-statistic results for IND20. (A) Outgroup $f_3$ statistics for IND20, computed in the form $f_3(X, \text{IND20}; \text{Yoruba})$ and ranked by decreasing affinity across comparison populations. (B) $D$-statistics, computed as $D(X, \text{Karitiana}; \text{IND20}, \text{Yoruba})$ and ranked by decreasing $Z$-score. Positive values indicate excess allele sharing between IND20 and population $X$ relative to Karitiana, while negative values indicate the reverse. Vertical dashed lines at $Z = \pm 3$ mark the conventional thresholds used to assess significance.

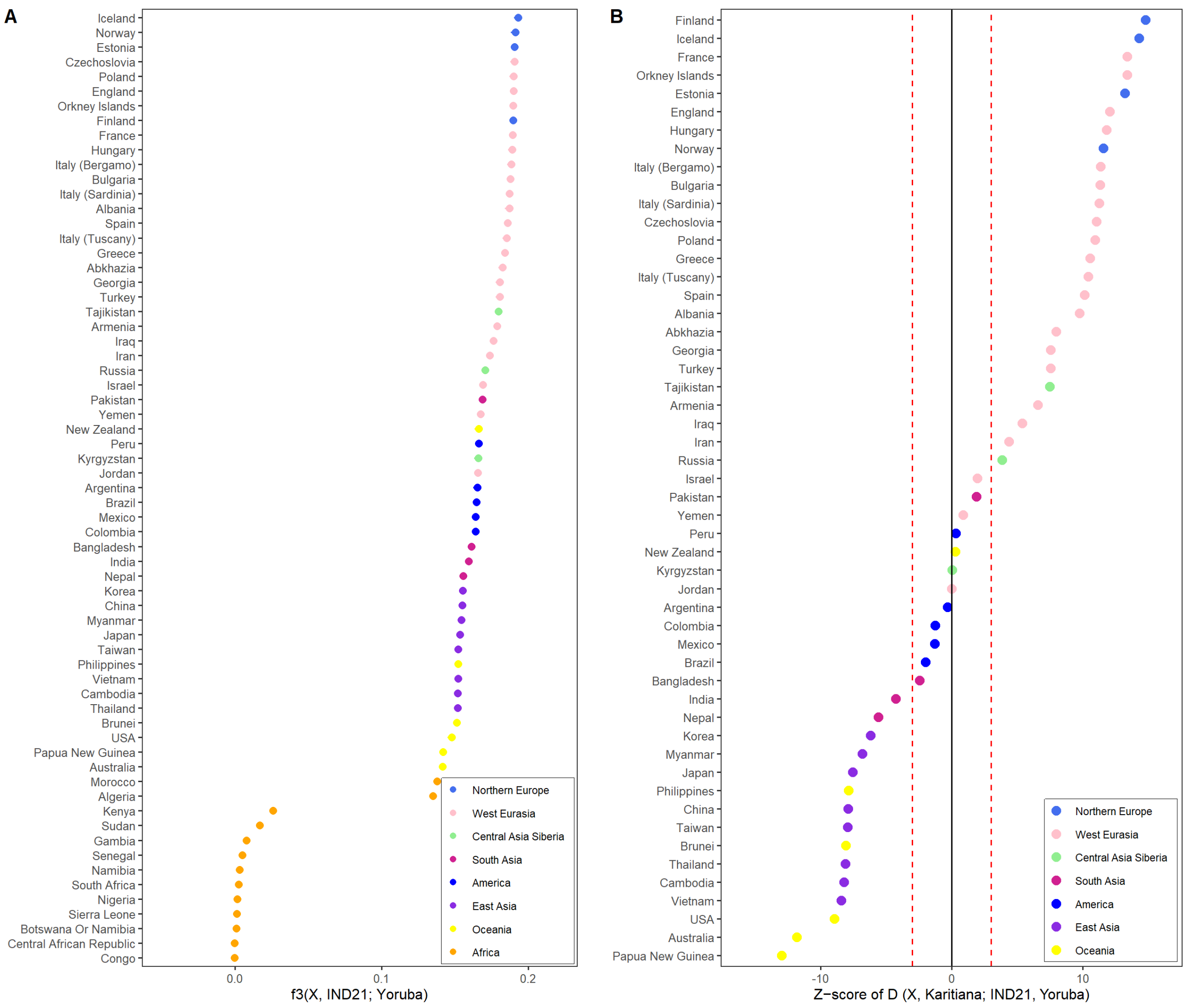


Figure S6: Outgroup $f_3$ and $D$-statistic results for IND21. (A) Outgroup $f_3$ statistics for IND21, computed in the form $f_3(X, \text{IND21}; \text{Yoruba})$ and ranked by decreasing affinity across comparison populations. (B) $D$-statistics, computed as $D(X, \text{Karitiana}; \text{IND21}, \text{Yoruba})$ and ranked by decreasing $Z$-score. Positive values indicate excess allele sharing between IND21 and population $X$ relative to Karitiana, while negative values indicate the reverse. Vertical dashed lines at $Z = \pm 3$ mark the conventional thresholds used to assess significance.

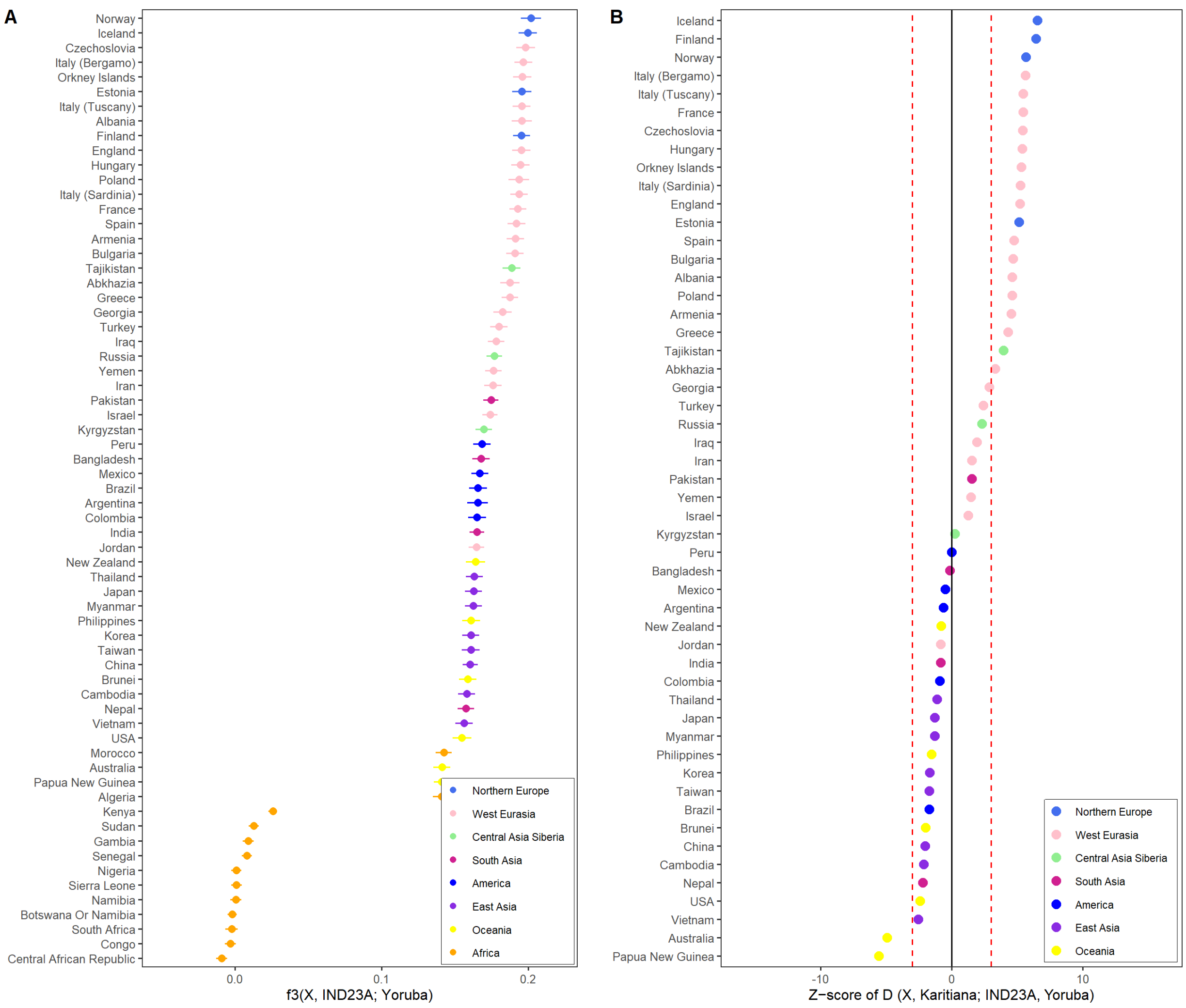


Figure S7: Outgroup $f_3$ and $D$-statistic results for IND23A. (A) Outgroup $f_3$ statistics for IND23A, computed in the form $f_3(X, \text{IND23A}; \text{Yoruba})$ and ranked by decreasing affinity across comparison populations. (B) $D$-statistics, computed as $D(X, \text{Karitiana}; \text{IND23A}, \text{Yoruba})$ and ranked by decreasing $Z$-score. Positive values indicate excess allele sharing between IND23A and population $X$ relative to Karitiana, while negative values indicate the reverse. Vertical dashed lines at $Z = \pm 3$ mark the conventional thresholds used to assess significance.

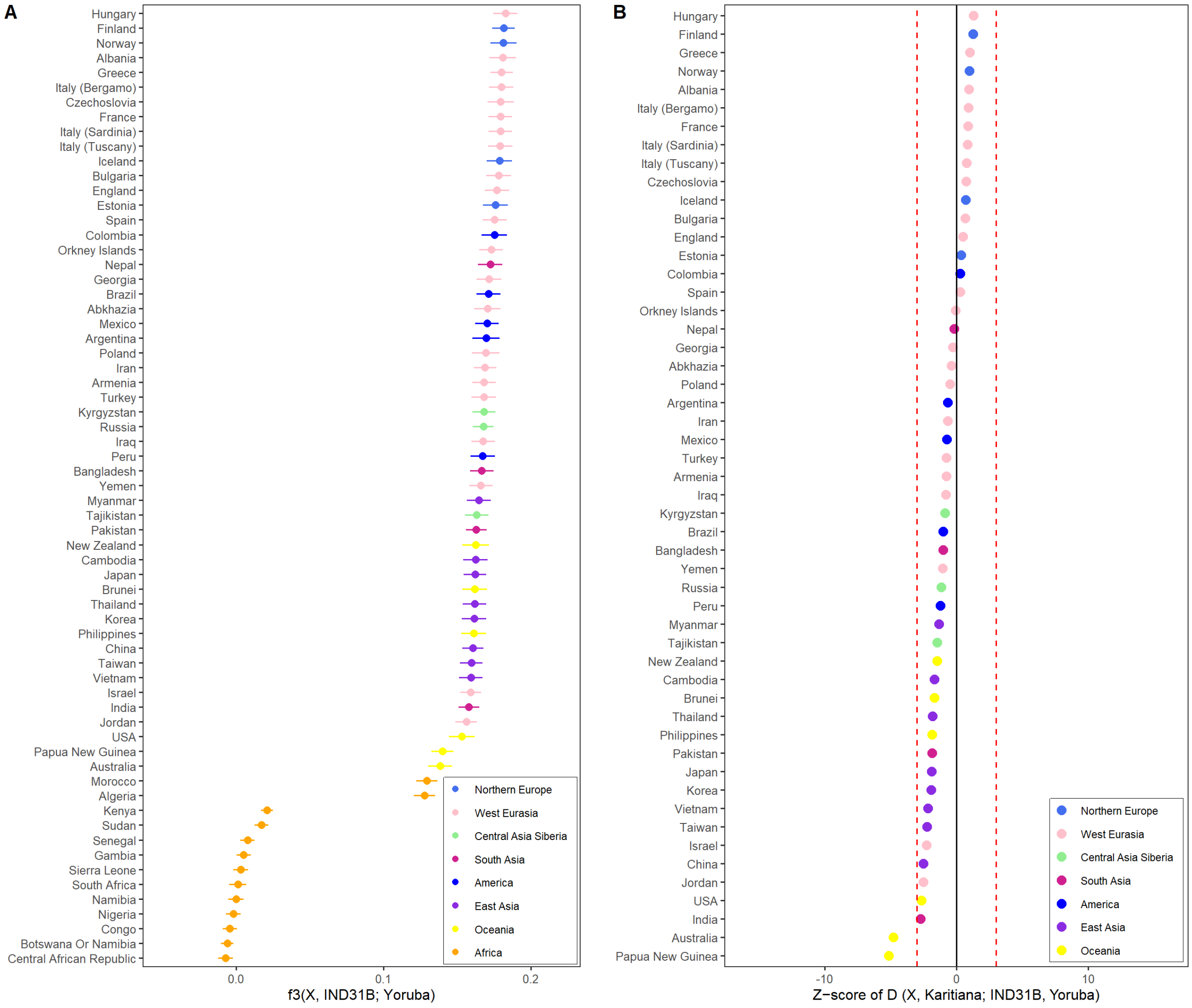


Figure S8: Outgroup $f_3$ and $D$-statistic results for IND31B. (A) Outgroup $f_3$ statistics for IND31B, computed in the form $f_3(X, \text{IND31B}; \text{Yoruba})$ and ranked by decreasing affinity across comparison populations. (B) $D$-statistics, computed as $D(X, \text{Karitiana}; \text{IND31B}, \text{Yoruba})$ and ranked by decreasing $Z$-score. Positive values indicate excess allele sharing between IND31B and population $X$ relative to Karitiana, while negative values indicate the reverse. Vertical dashed lines at $Z = \pm 3$ mark the conventional thresholds used to assess significance.

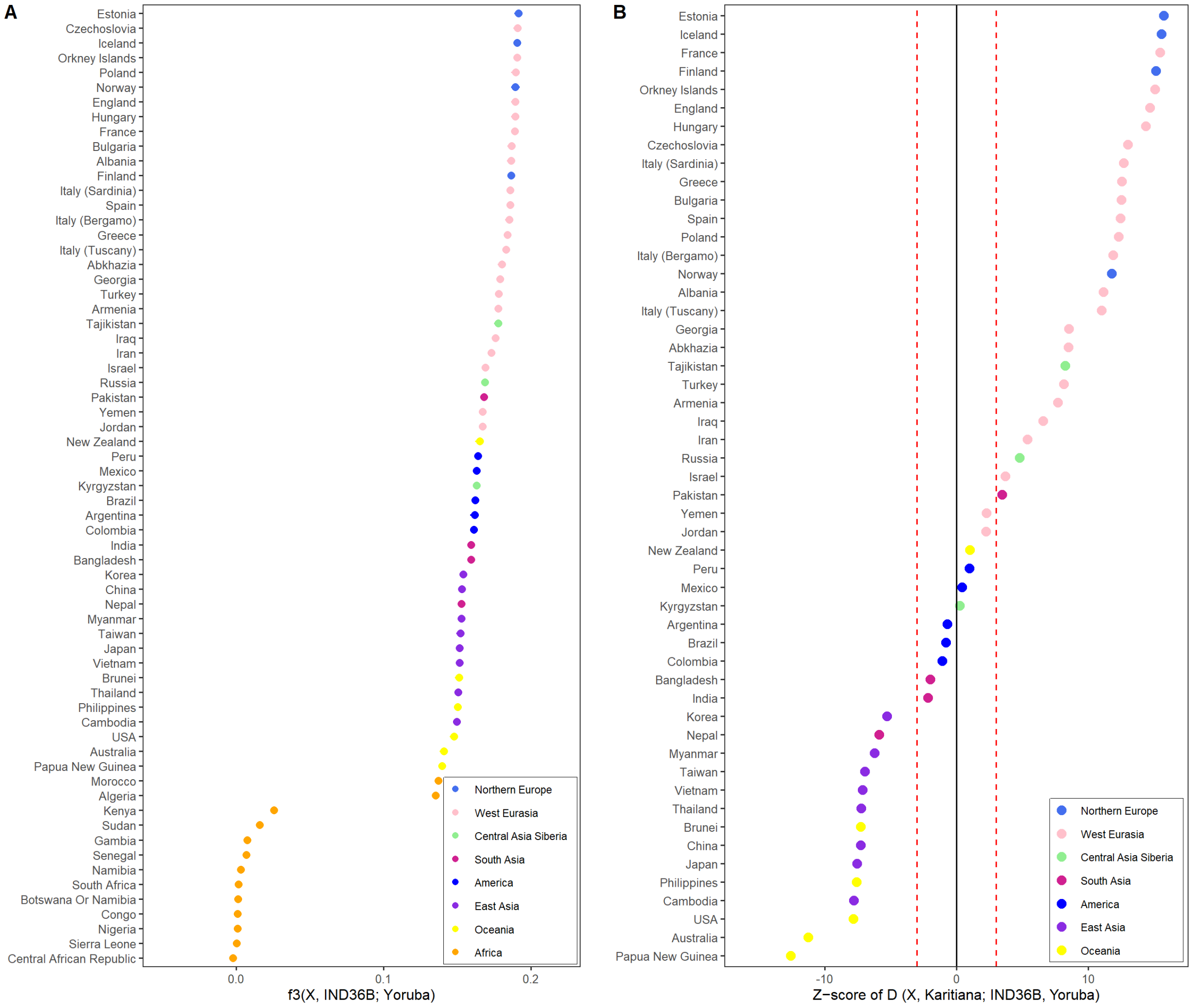


Figure S9: Outgroup $f_3$ and $D$-statistic results for IND36B. (A) Outgroup $f_3$ statistics for IND36B, computed in the form $f_3(X, \text{IND36B}; \text{Yoruba})$ and ranked by decreasing affinity across comparison populations. (B) $D$-statistics, computed as $D(X, \text{Karitiana}; \text{IND36B}, \text{Yoruba})$ and ranked by decreasing $Z$-score. Positive values indicate excess allele sharing between IND36B and population $X$ relative to Karitiana, while negative values indicate the reverse. Vertical dashed lines at $Z = \pm 3$ mark the conventional thresholds used to assess significance.

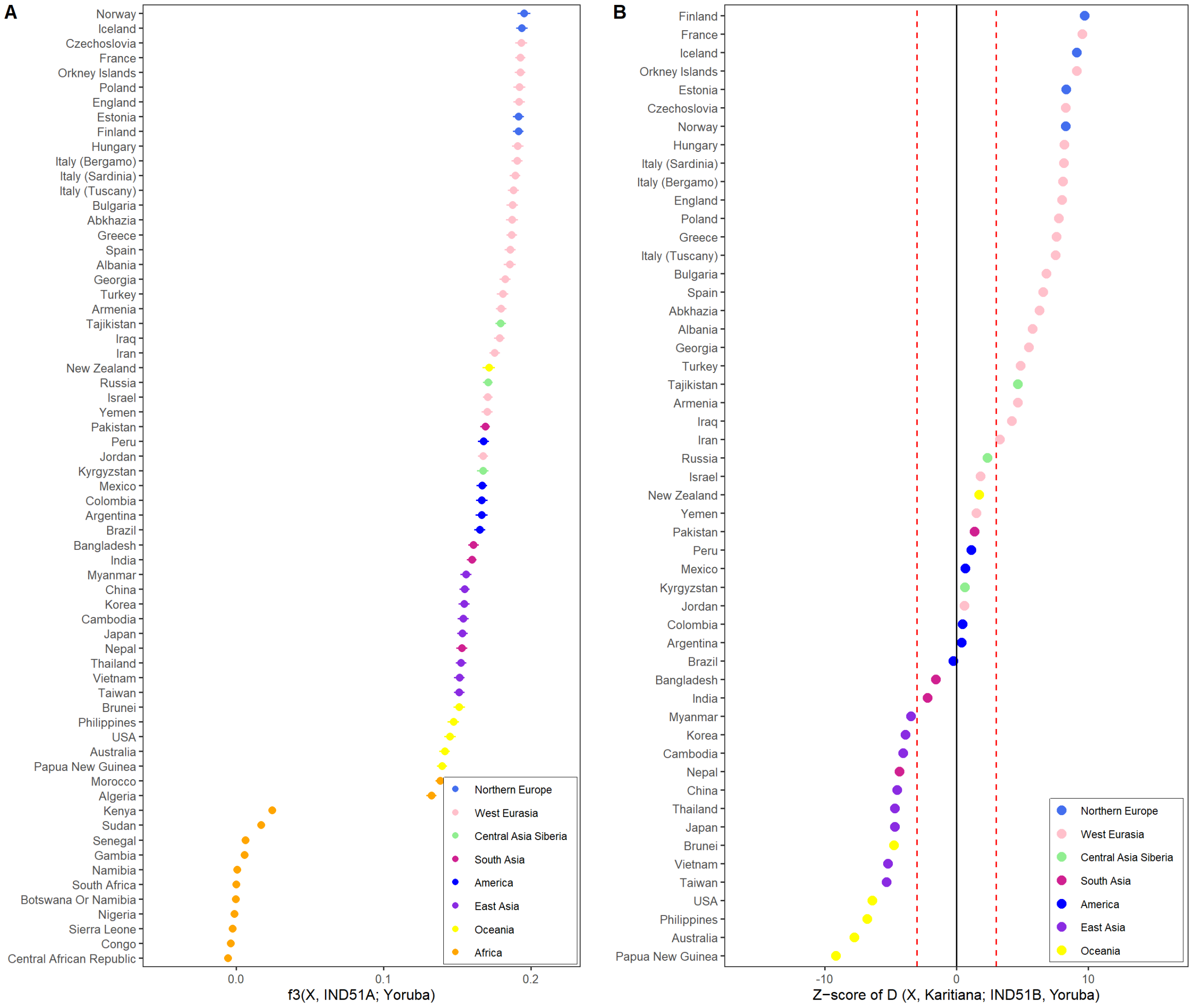


Figure S10: Outgroup $f_3$ and $D$-statistic results for IND51A. (A) Outgroup $f_3$ statistics for IND51A, computed in the form $f_3(X, \text{IND51A}; \text{Yoruba})$ and ranked by decreasing affinity across comparison populations. (B) $D$-statistics, computed as $D(X, \text{Karitiana}; \text{IND51A}, \text{Yoruba})$ and ranked by decreasing $Z$-score. Positive values indicate excess allele sharing between IND51A and population $X$ relative to Karitiana, while negative values indicate the reverse. Vertical dashed lines at $Z = \pm 3$ mark the conventional thresholds used to assess significance.

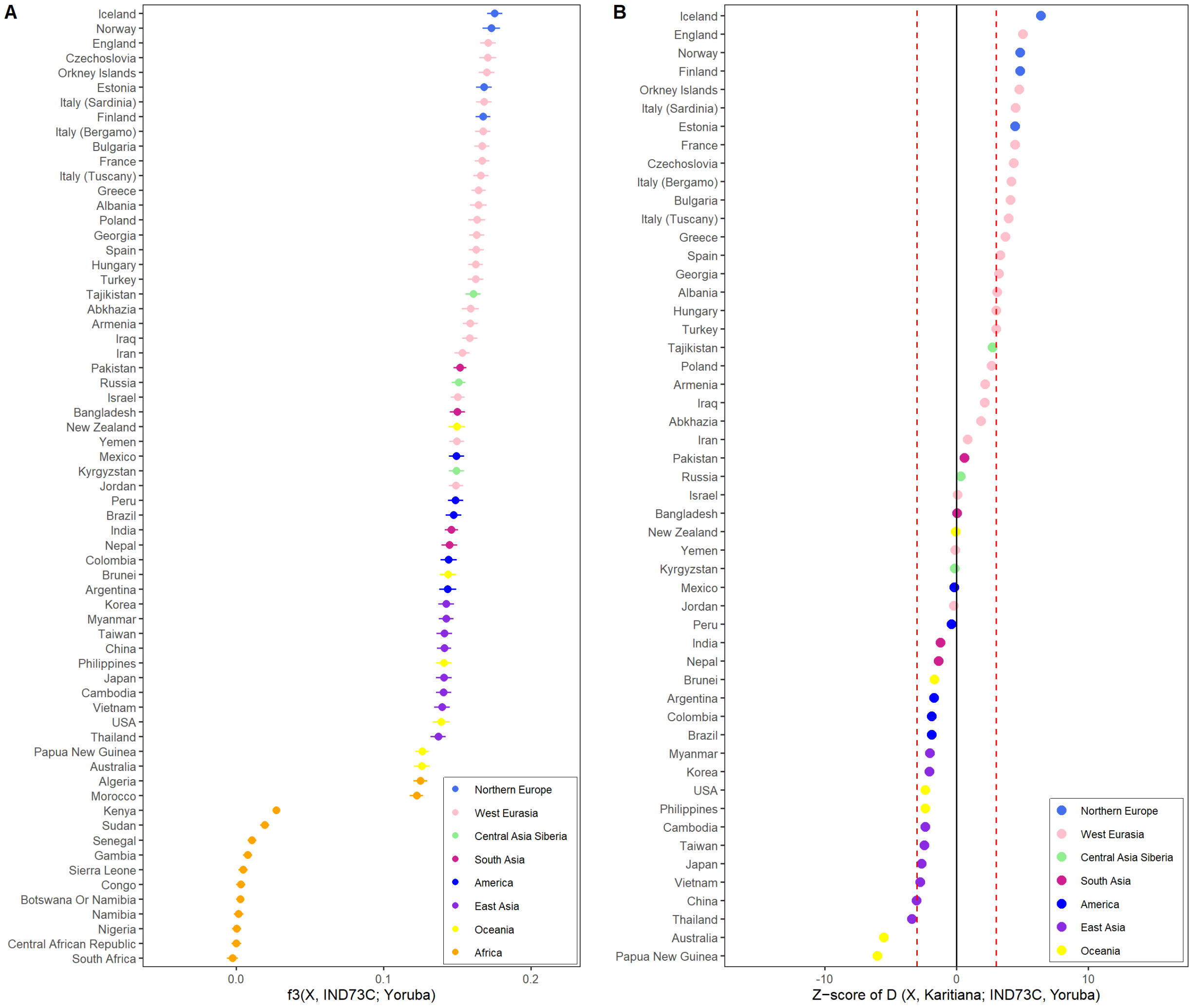


Figure S11: Outgroup $f_3$ and $D$-statistic results for IND73C. (A) Outgroup $f_3$ statistics for IND73C, computed in the form $f_3(X, \text{IND73C}; \text{Yoruba})$ and ranked by decreasing affinity across comparison populations. (B) $D$-statistics, computed as $D(X, \text{Karitiana}; \text{IND73C}, \text{Yoruba})$ and ranked by decreasing $Z$-score. Positive values indicate excess allele sharing between IND73C and population $X$ relative to Karitiana, while negative values indicate the reverse. Vertical dashed lines at $Z = \pm 3$ mark the conventional thresholds used to assess significance.

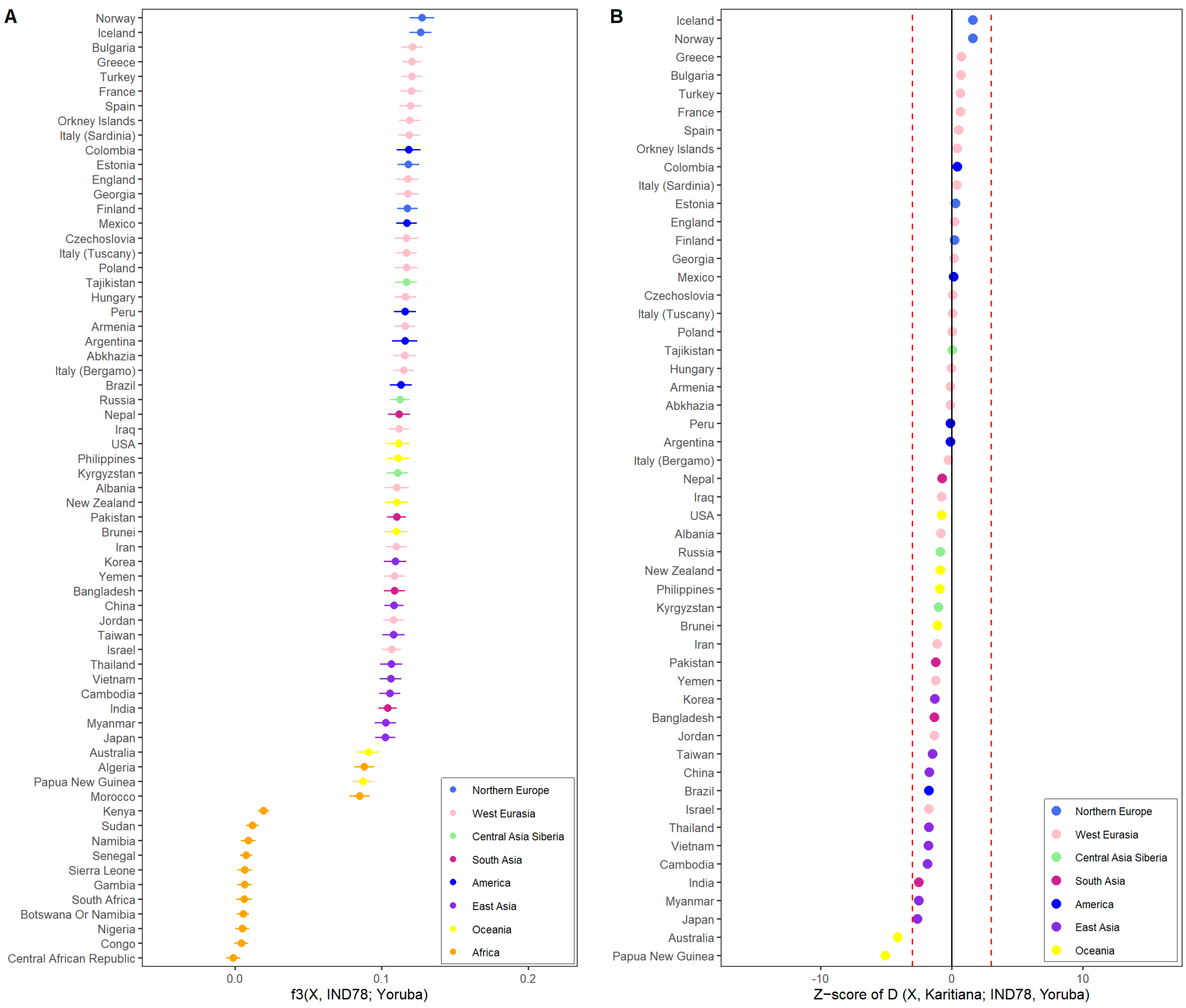


Figure S12: Outgroup $f_3$ and $D$-statistic results for IND78. (A) Outgroup $f_3$ statistics for IND78, computed in the form $f_3(X, \text{IND78}; \text{Yoruba})$ and ranked by decreasing affinity across comparison populations. (B) $D$-statistics, computed as $D(X, \text{Karitiana}; \text{IND78}, \text{Yoruba})$ and ranked by decreasing $Z$-score. Positive values indicate excess allele sharing between IND78 and population $X$ relative to Karitiana, while negative values indicate the reverse. Vertical dashed lines at $Z = \pm 3$ mark the conventional thresholds used to assess significance.

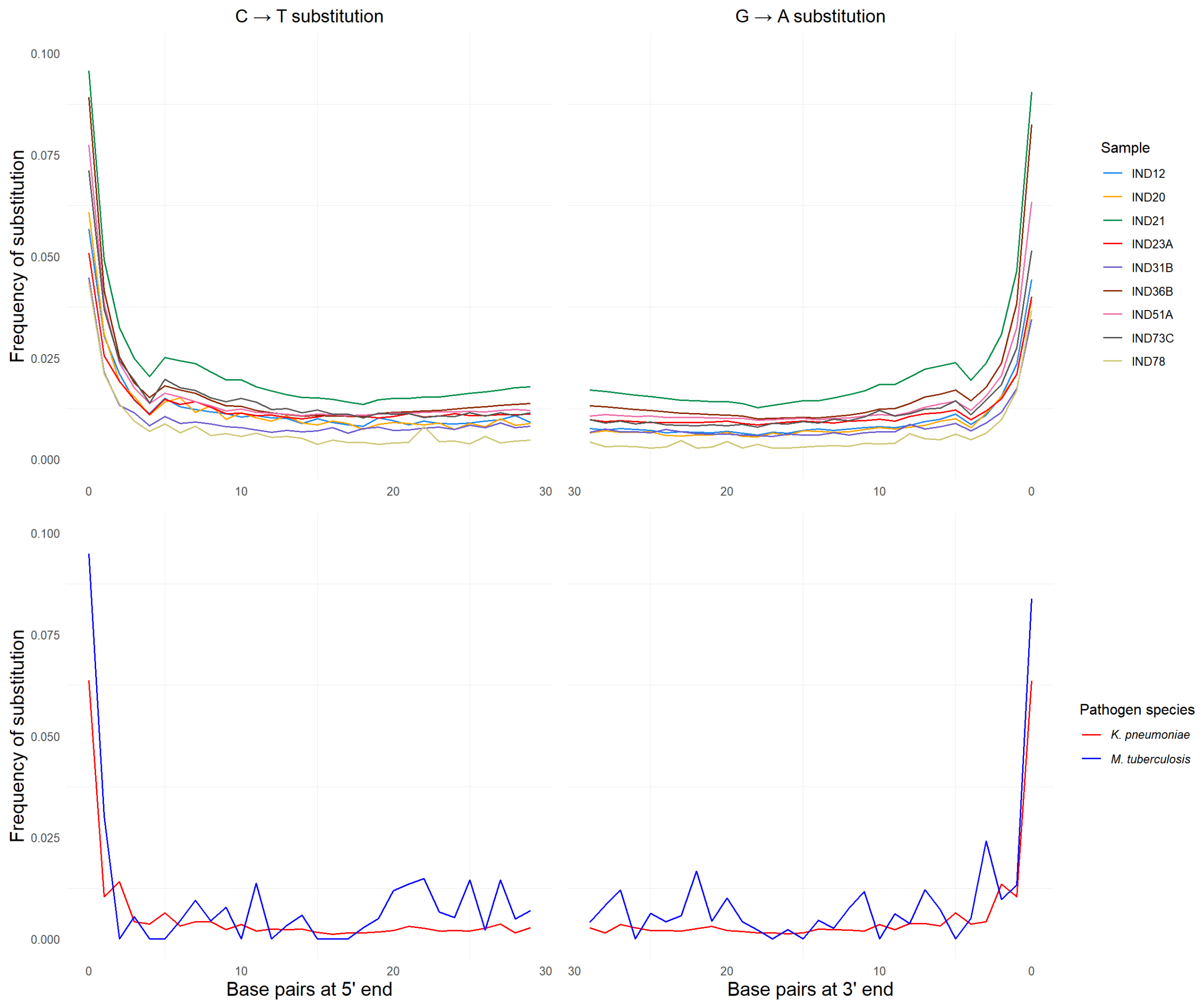


Figure S13: Postmortem damage profiles inferred using PMDtools. Left panels show 5′-end C→T substitution frequencies, and right panels show 3′-end G→A substitution frequencies across read positions. Damage patterns are displayed for the nine Pilar individuals (top) and for *K. pneumoniae* and *M. tuberculosis* reads recovered from IND21, the individual yielding the highest number of pathogen-derived sequences (bottom). Elevated terminal deamination rates in both human and microbial DNA are consistent with authentic ancient DNA. The *M. tuberculosis* curve exhibits greater variance due to its substantially lower read depth relative to *K. pneumoniae*.

Table S1: Uniparental haplogroups and autosomal coverage metrics for the nine Pilar individuals.

| Sample | mtDNA haplogroup | Y haplogroup | Mean depth | Mean coverage |
|---|---|---|---|---|
| IND12 | H1u | P-P337 | 0.321 | 0.054 |
| IND20 | W1c | R-Z159 | 0.347 | 0.051 |
| IND21 | H1b | R-S23634 | 1.578 | 0.486 |
| IND23A | H70 | A-Y156117 | 0.450 | 0.084 |
| IND31B | H26a1 | R-YP5802 | 0.264 | 0.051 |
| IND36B | H2a2 | R-Y130457 | 2.028 | 0.590 |
| IND51A | U2e2a1c | R-L52 | 1.167 | 0.201 |
| IND73C | I1a1 | R-Y3159 | 0.771 | 0.115 |
| IND78 | H3+152 | R-M269 | 0.295 | 0.047 |
